## Supplemental Figure for "Idiosyncratic Perception: A Link Between Acuity, Perceived Position, and Apparent Size"

### Supplemental Figures

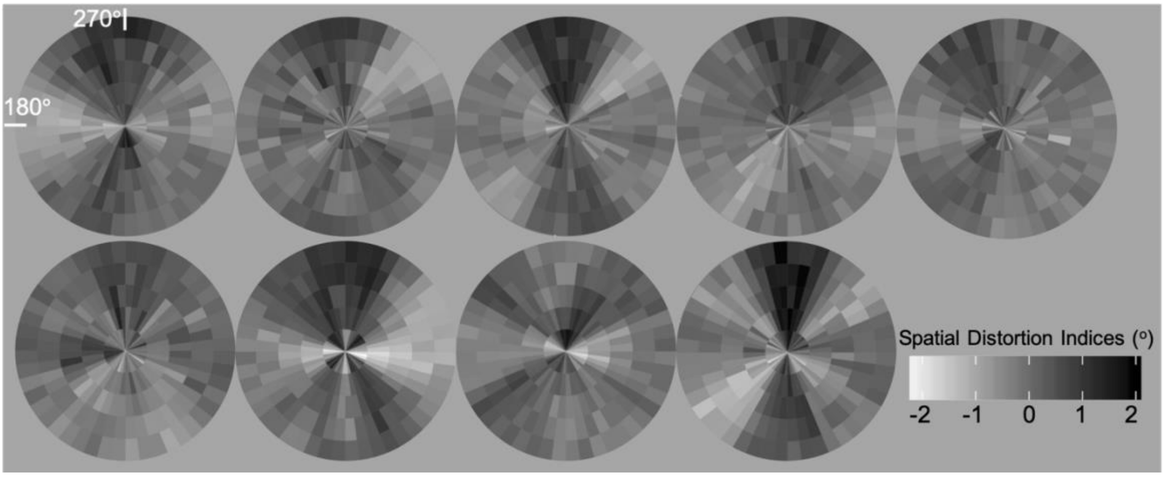


***Figure S1.*** The gray-scale version of the spatial distortion maps reported in Experiment 1. Brighter color (negative spatial distortion indices) indicates contraction of visual space and darker color (positive spatial distortion indices) represents expanded visual space.

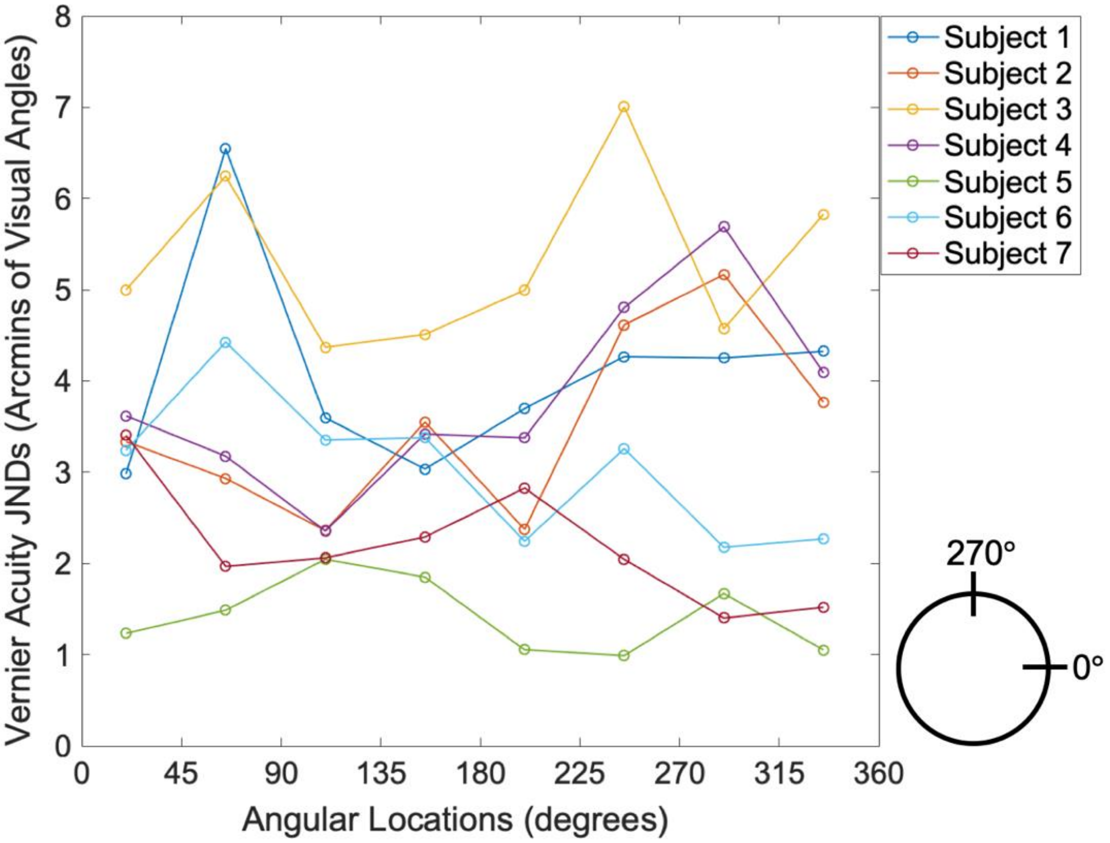


***Figure S2.*** Change of Vernier acuity as a function of the angular locations tested for every observer. Subject 1 and Subject 4 are authors. The layout of the angular locations is shown on the bottom right corner.

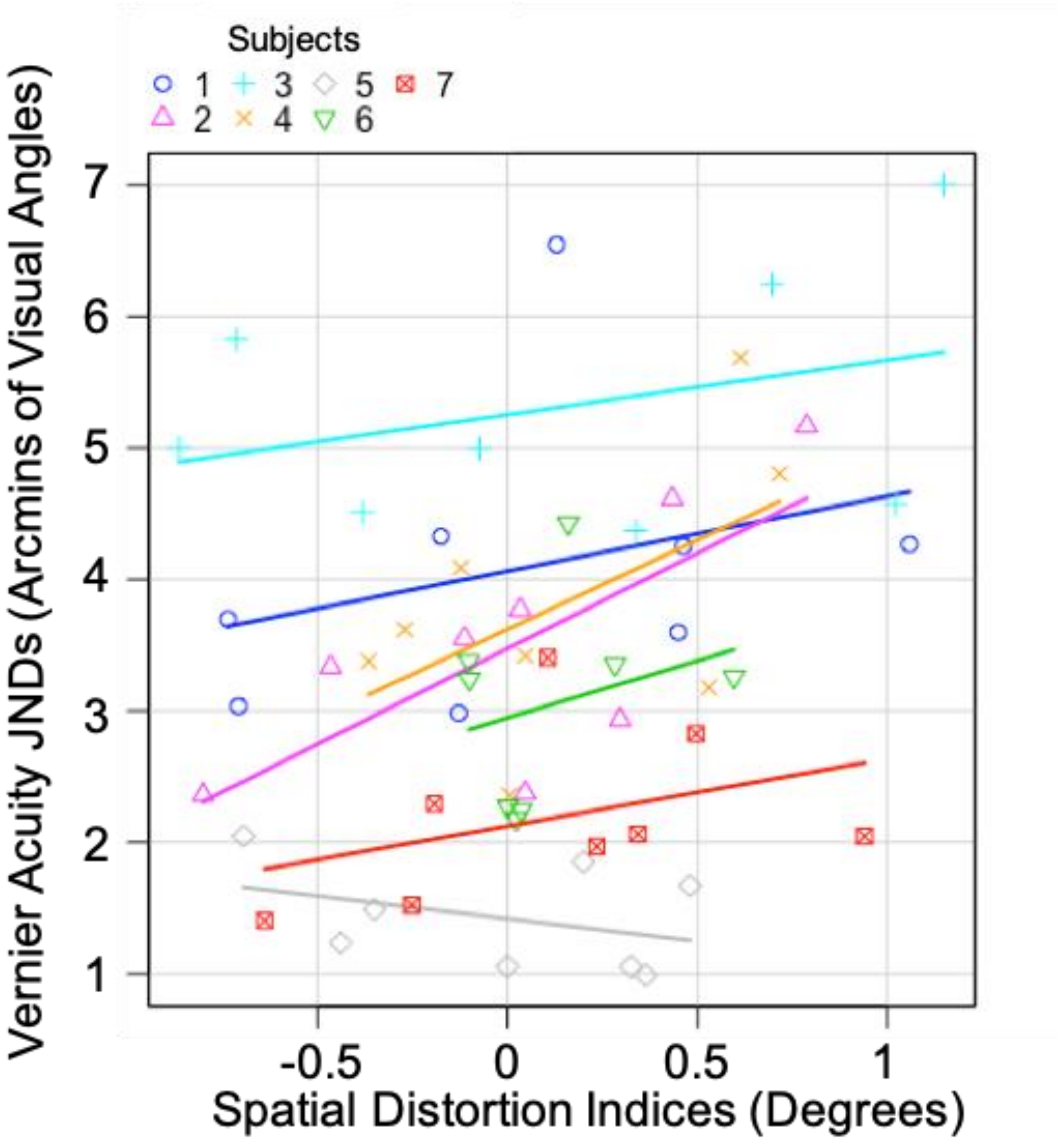


***Figure S3.*** Correlation between spatial distortion indices and Vernier acuity JNDs for each observer. Each observer had 8 pairs of data, corresponding to 8 angular locations tested in Experiment 2. Different symbols represent different observers. Lines are regression lines fitted based on each observer's data. The Pearson's correlations for individual subjects were 0.31, 0.73, 0.34, 0.55, -0.37, 0.26, 0.38 (listed in the same order as the figure legend) and the mean correlation calculated from Fisher transformation was 0.34. Note that the only observer who did not show the same trend (displayed as gray diamond) had the smallest JNDs (i.e., best acuity), so we speculated that it might be subject to a ceiling effect. This could affect the measured variability of Vernier acuity across different locations and thus influence the correlation calculated based on it.

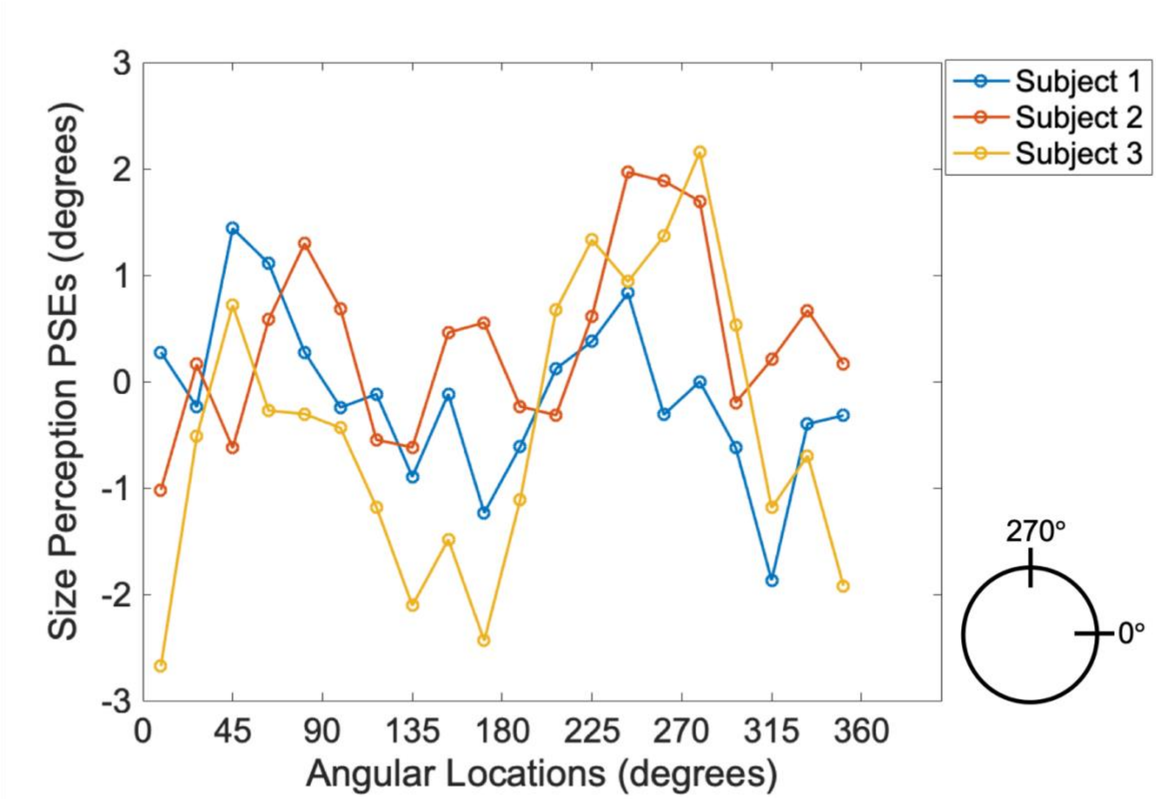


***Figure S4.*** The change of perceived size of the arc stimuli as a function of the angular locations tested for every observer. Subject 1 and Subject 2 are authors. The layout of the angular locations is shown on the bottom right corner.

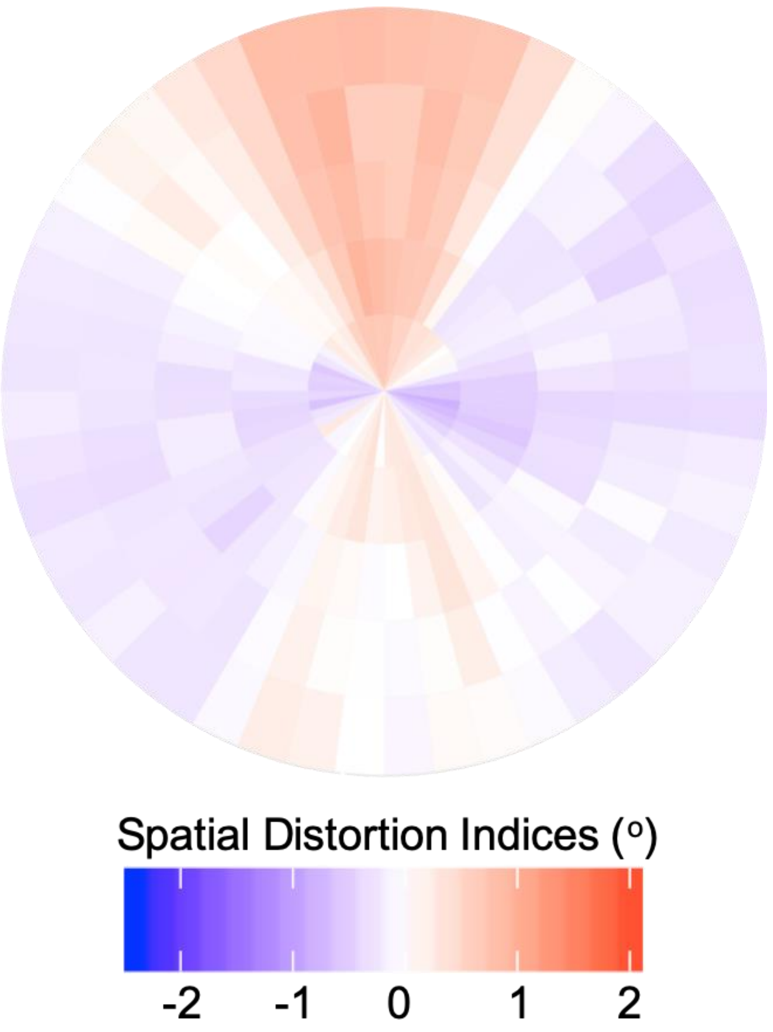


***Figure S5***. The group mean of the spatial distortion pattern calculated from Experiment 1 ($N = 9$). Blue area represents visual space compression and red represents expansion. The group average effect is notable and also comports with previous findings on various spatial biases or spatial anisotropies of visual performance (e.g., Abrams, Nizam, & Carrasco, 2012; Low, 1943), but it is smaller and easily washed out by the idiosyncratic differences found at the individual-subject level (Fig. 1b).
